# StructureSAFE: A structure-aware chemical language model for unified hit identification and lead optimization

**DOI:** 10.64898/2026.06.28.735128

**Authors:** Bo Yang, Ke Xu, Chijian Xiang, Byeongtak Lee, Yunong Xu, Tongtong Li, Yutong Shi, Anton V. Sinitskiy, Jianing Li

**Affiliations:** Borch Department of Medicinal Chemistry and Molecular Pharmacology, Purdue University, West Lafayette, IN 47907, USA; Department of Horticulture & Landscape Architecture, Purdue University, West Lafayette, IN 47907, USA; ML LC, Southborough, MA 01772, USA; College of Professional Studies, Northeastern University, Boston, MA 02115, USA

**Keywords:** Structure-based drug design, Molecular generation, Chemical language model, Lead optimization, *De novo* hit identification, SAFE representation

## Abstract

Structure-based generative models (SBGMs) hold great promises for accelerating drug discovery by enabling target-aware molecular design. However, existing approaches face fundamental challenges: three-dimensional graph-based models can explicitly incorporate protein structural information but often generate chemically implausible molecules due to limited training data, while chemical language models (CLMs) produce chemically plausible molecules but struggle to effectively leverage three-dimensional structural information for structure-conditioned generation and hard to incorporate lead optimization functionality due to the nature of SMILES string. Here, we present StructureSAFE, a structure-aware chemical language model that resolves this trade-off by integrating protein structural and evolutionary encoders with the SAFE molecular representation via pretraining and finetuning training scheme, enabling both de novo hit identification and a comprehensive suite of lead optimization subtasks within a unified framework. Comprehensive benchmarking on the MolGenBench dataset demonstrates that StructureSAFE achieves state-of-the-art (SOTA) performance across multiple metrics, with particularly pronounced improvements in chemical plausibility relative to graph-based models lacking pretraining. Evaluation on a rigorously constructed held-out test set further confirms its ability to generate drug-like, synthetically accessible molecules with competitive predicted binding affinities for previously unseen targets on both hit identification and lead optimization setting. In silico case studies across four therapeutically relevant targets validate its capacity to generate chemically plausible molecules that recapitulate key binding interactions of known high-affinity ligands while proposing novel interactions for potential better affinity and exploring previously unknown regions of chemical space. Taking together, StructureSAFE represents a versatile and practical tool to provide high-quality candidate molecules for augmenting medicinal chemistry workflows in both hit identification and lead optimization campaigns.

## Introduction

Drug discovery is a costly, time-consuming and failure-prone process, where the total cost per approved drug is more than one billion dollars.^1^ While clinical phase is the most expensive phase in drug discovery pipeline, the sequential nature and highly cumulative failure rates indicate that the majority cost of approved drug comes from accumulated failures in the preclinical discovery phase.^1^ In the highly complex, multi-parameter optimization preclinical discovery phase, acquiring molecules with optimized affinity on both targets and anti-targets plays significant role which could provide benefits on other aspects of preclinical drug discovery.^2^ Thus, methods help to find high affinity ligands for given receptors hold the potential to largely speed up the complex multi-parameter optimization and reduce the time and cost of the drug discovery process.

Structure-based drug discovery (SBDD) encompasses a range of widely used methods for identifying high-affinity hits and optimizing ligand affinity from binding pocket structures.^3^ Recently, as the development of biophysical techniques and structural predictive tools like Alphafold3^4^ and RosettaFold^5^, the structural information becomes more and more accessible to researchers. Thus, SBDD plays an even more important role in optimizing the binding affinities of ligands. Besides traditional SBDD methods like docking and molecular dynamics (MD) simulation, with the development of deep learning, structure-based generative models (SBGMs) have started to appear and soon become handy tools for drug discovery specialists.^6-8^ Utilizing more accessible structural information, they could directly *de novo* generate molecules or optimize known molecules to give potential high-affinity compounds, as well as explore novel chemical spaces, holding great potential to speed up drug discovery processes at a lower cost.

Current SBGMs are mostly diffusion models and flow models that operate in 3D space with point cloud representation of both protein and ligand atoms.^9-11^ Although they could explicitly utilize 3D special information as conditions and restrictions to generate ligands with 3D conformation, lots of generated molecules are chemically implausible, which could be attributed to suboptimized learning of chemical prior information due to limited protein-ligand data available.^12^ Meanwhile, chemical language models (CLM) perform well on generating chemically plausible molecules with significant efficiency by pretraining on a large scale of molecular data and finetuning with different substream tasks.^13-15^ However, the ability of them to do structural-based generation is limited compared to 3D graph-based models due to the implicit embedding of 3D protein structural information.^16^ Meanwhile, since majority of CLMs work on SMILES representation, it is relatively inconvenient to do lead optimization related tasks due to the inherit disadvantages of SMILES: It could not provide a contiguous representation of a molecular substructure.^17^ In result, the majority of lead optimization tasks on binding affinity from CLM are done by incorporating reinforcement learning techniques that utilize Vina docking scores as rewards to guide the generation.^18^ Comparing to directly optimize molecules based on structural information, docking score would inevitably bring its own bias and mislead the optimization. Recently developed SAFE representation could help to solve this problem due to its ability of represent molecules as fragment-wise strings, which enables its smooth incorporation for all subtasks in lead optimization.^17, 19^

To solve the problem mentioned above, we developed StructureSAFE, a structure-aware chemical language model capable of both hit identification and lead optimization. StructureSAFE is pretrained on 1.5B molecules from ZINC22 to learn the underlying chemical syntax and grammar of SAFE strings. This pretraining strategy enables us to leverage the broad generative capabilities inherent to the SAFE representation, supporting both de novo molecular generation and a diverse set of lead optimization subtasks (including linker design, scaffold hopping, fragment growing, scaffold decoration, and fragment elaboration) while preserving the chemical plausibility of generated molecules (Figure 1).^17^ The pretrained model is subsequently finetuned on ~0.7 M protein-ligand complex data to enable target-specific molecular generation conditioned on given protein binding pockets.^13, 20^ By training on one of the most comprehensive 3D protein–ligand complex datasets available, and incorporating information-rich 3D protein structural features, evolutionary information, and optionally ligand and protein–ligand interaction information, StructureSAFE is designed to maximize generalizability and performance in structure-conditioned molecular generation. In our comprehensive benchmarking experiments, StructureSAFE achieves state-of-the-art performance across multiple metrics on the MolGenBench dataset for de novo design, with particularly strong results in chemical plausibility and hit rediscovery. Evaluated on a carefully constructed held-out test set, StructureSAFE consistently generates drug-like molecules with competitive Vina docking scores for previously unseen targets. In lead optimization tasks, StructureSAFE demonstrates competitive performance across multiple subtasks, underscoring its versatility for both de novo hit identification and structure-guided lead refinement. Finally, in silico case studies further validate its capacity to design novel drug-like binders and refine known ligands through scaffold hopping and fragment elaboration on target protein pockets, while expanding into previously unexplored regions of chemical space.

**Figure 1.**
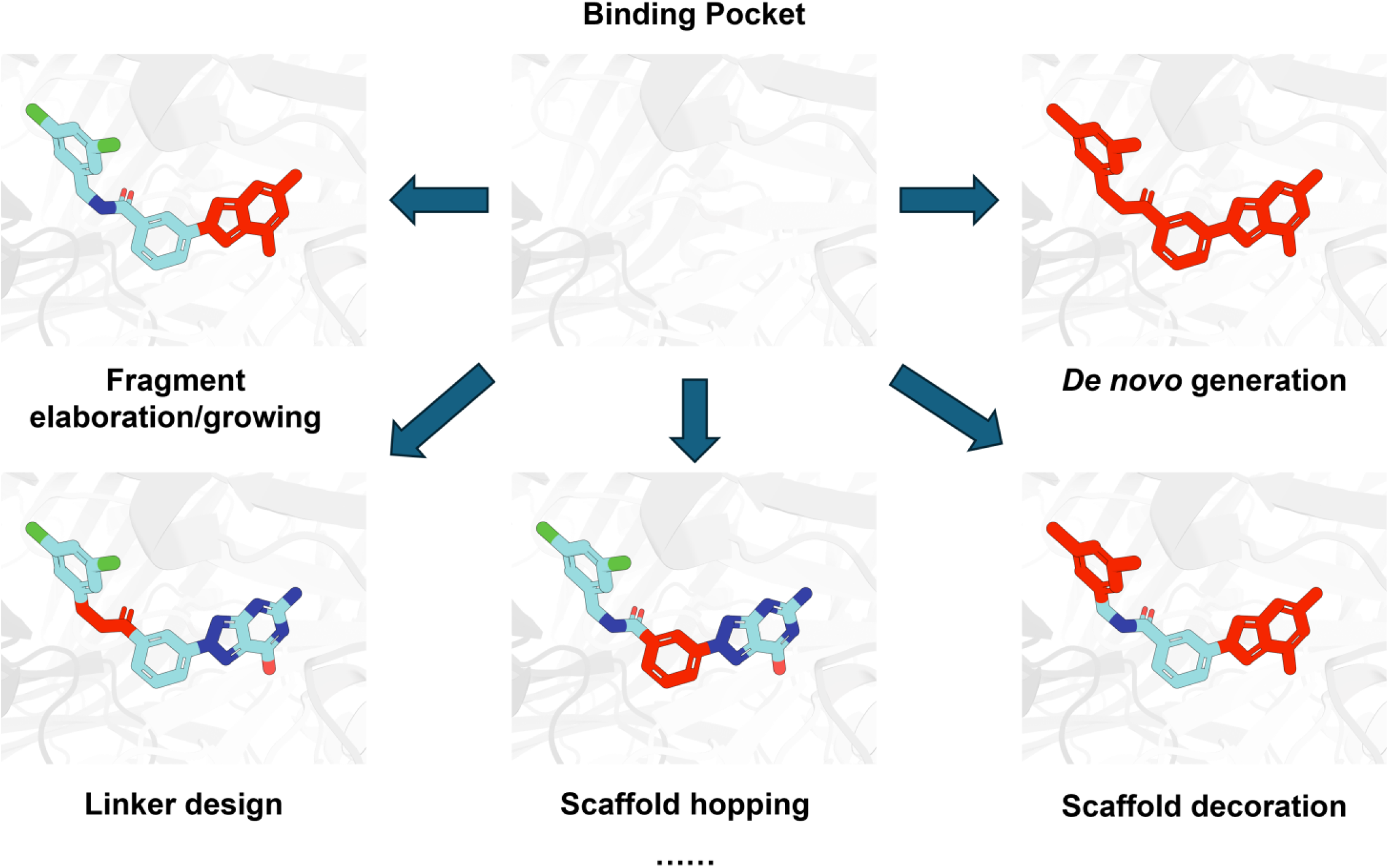
Structure-based drug design tasks could be performed by StructureSAFE. Most CLMs with SMILES as representation could only perform *De novo* generation task.

**Figure 1.**
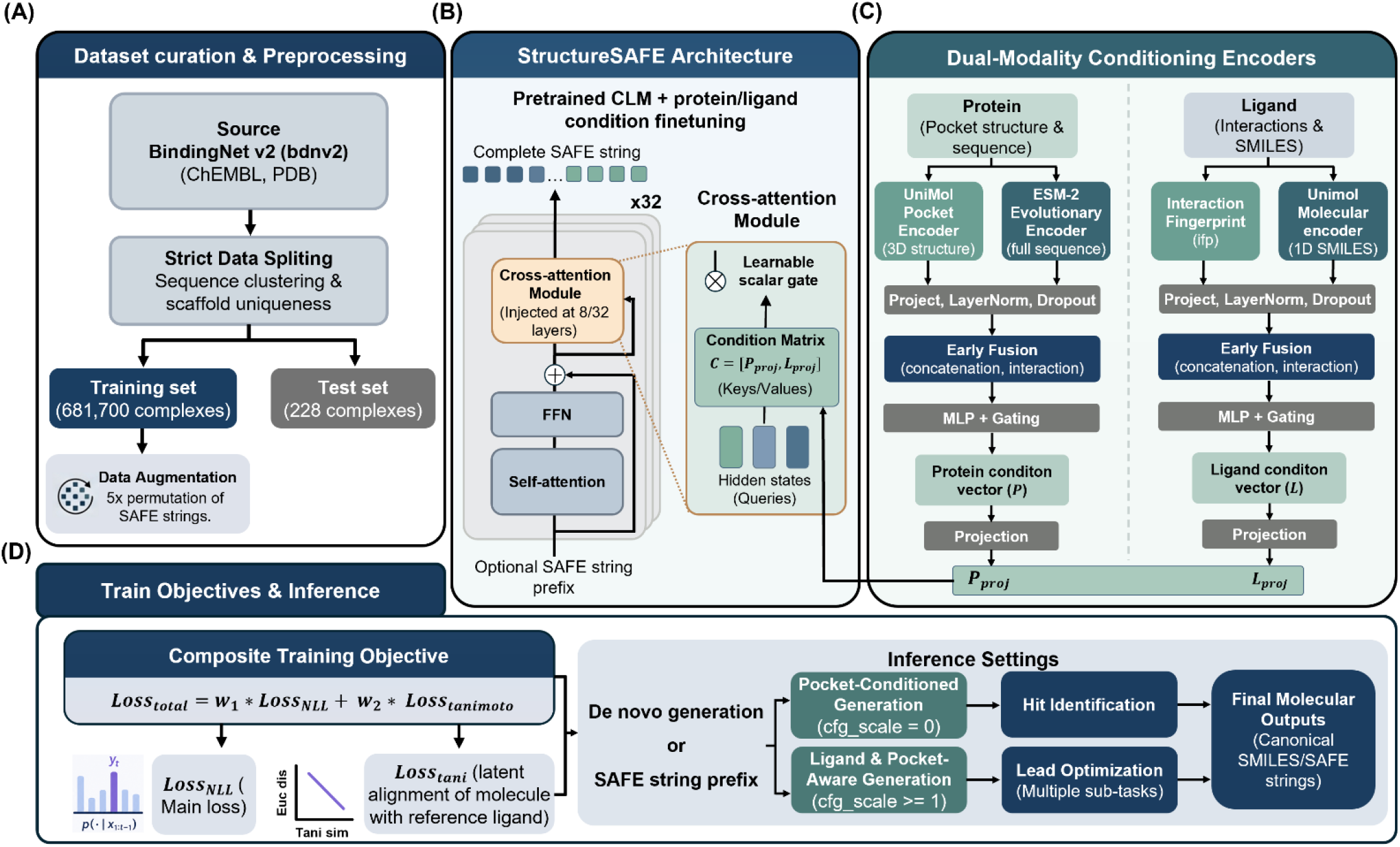
Overview of StructureSAFE. (A) Data curation pipeline. Training data was augmented five-fold through SAFE string randomization to enhance the model’s capture of chemical prior information and mitigate overfitting. (B) StructureSAFE architecture. After pretraining, the backbone model undergoes full fine-tuning with conditioning information incorporated as keys and values in the cross-attention module, enabling structure-conditioned molecular generation. (C) Conditioning module. StructureSAFE integrates four sources of conditioning information including pocket three-dimensional structural features (UniMol), pocket evolutionary information (ESM-2), and optionally protein–ligand interaction fingerprints (PLEC) and known ligand embeddings, and conditions the generation process via cross-attention. (D) Training objectives and inference workflow. Model training jointly optimizes negative log-likelihood (NLL) loss and Tanimoto similarity loss. At inference, the trained StructureSAFE model supports both de novo hit identification and lead optimization subtasks, conditioned on protein structural information and/or known ligand information.

## Method

### Dataset

CrossDocked2020 is a widely applied dataset used for structure-based generative model training, which contains 100,000 protein-ligand complexes as training set and 100 protein-ligand complexes as test set based on protein sequence similarity-based splitting. However, its uniqueness of ligands and quality of protein-ligand binding has been raising concerns about using it as training set. In this study, we utilized BindingNet v2 (bdnv2) dataset as our data source.^20^ Bdnv2 contains 689,796 protein-ligand (PL) complexes extracted or modeled from Protein Data Bank (PDB) and ChEMBL databases, representing one of the richest 3D protein-ligand data sources. To test generalization ability of our model, we applied strict data-splitting based on both protein sequence and ligand scaffolds. In detail, we did sequence-based clustering using mmseqs2 with 30% sequence identify as cut-off. After clustering, we took all singlet proteins and only left complexes with highest binding affinity known ligand (measure by bioassay and provided by bdnv2). Then, we standardized the SMILES of those ligands. BM scaffolds of those ligands as well as ligands from non-singlet proteins were extracted, only complexes made of unique BM scaffolds ligands and singlet proteins were selected as test set candidates. Finally, to avoid taking too many complexes from training set, we manually inspected the test set candidates and took proteins with less than 200 known ligands and their highest affinity known ligands as final test set. And all other proteins and their known ligands complexes are grouped as training sets. After splitting, training set and test set contain 681,700 and 228 protein-ligand complexes, respectively. Ligands are represented as standardized SMILES and are converted to SAFE strings on-the-fly during training. This splitting ensures limited sequence similarity between proteins and no scaffold overlapping between ligands in training and test set, allowing reliable testing of model’s generalization ability.

To further boost the performance of our model, we have augmented the data by utilizing the permutation invariance characteristics of SAFE string. In detail, for each SAFE string of ligands in training set, we randomly switched the ordering of each fragment separated by “.” for five times, allowing 5 folds data augmentation on our training data (Figure 2A). Finally, to allow fair comparison with other SOTA models trained using CrossDocked2020 dataset. We have also finetuned our model using CrossDocked2020 with same data augmentation and comparing the results with other models on MolGenBench benchmarking datasets.^21^

**Figure 2.**
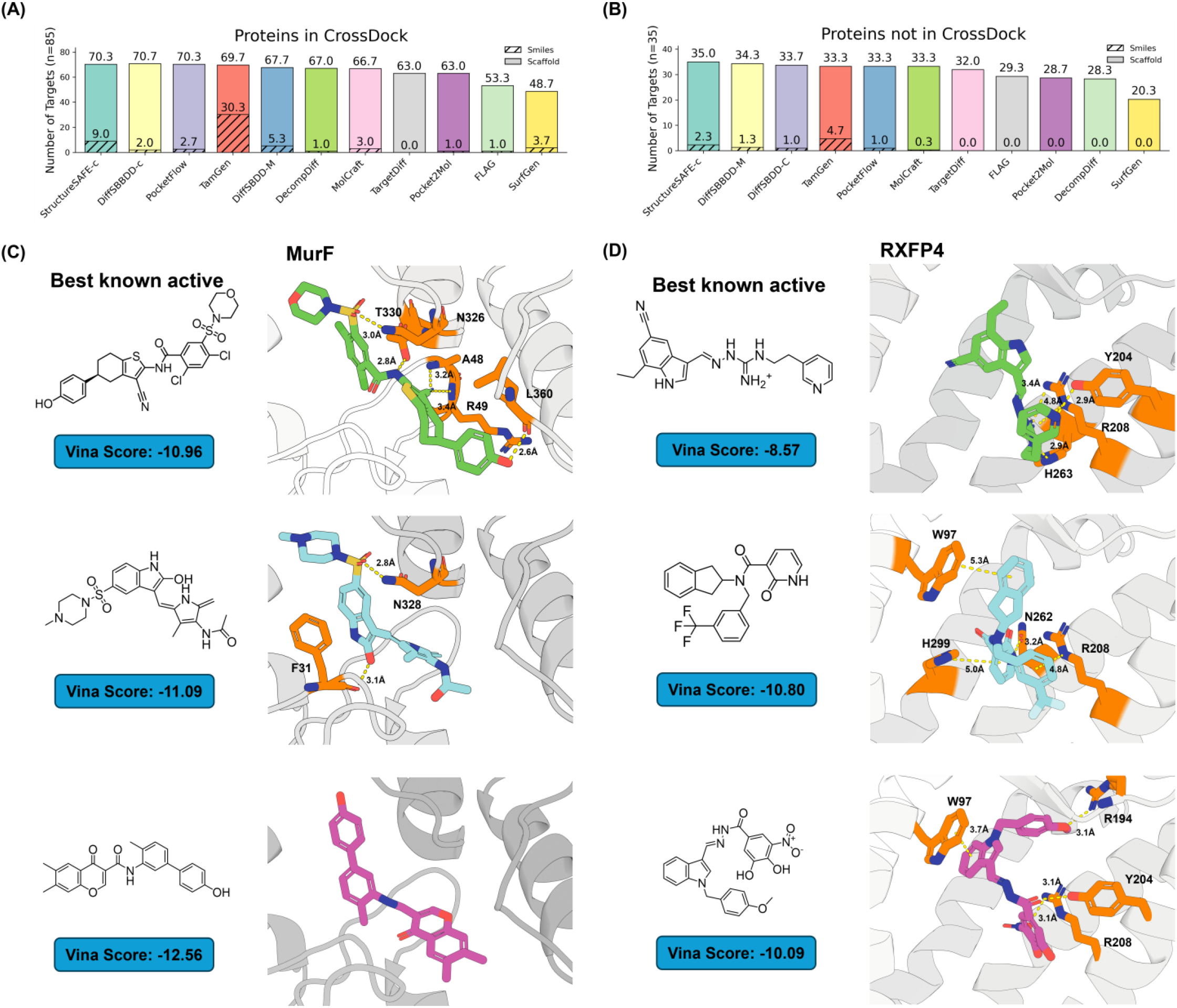
De novo hit identification performance of StructureSAFE on the MolGenBench dataset and in silico case studies. (A) StructureSAFE-c achieves second-ranked performance in regenerating known SMILES and first-ranked in regenerating known scaffolds for proteins in the MolGenBench dataset that are present in the CrossDocked2020 training set. (B) StructureSAFE-c achieves second-ranked performance in regenerating known SMILES and first-ranked performance in regenerating known scaffolds for proteins in the MolGenBench dataset that are absent from the CrossDocked2020 training set. (C, D) StructureSAFE was applied to generate de novo hit molecules conditioned on protein structure for the ligase target MurF (C) and the GPCR target RXFP4 (D). For each target, all generated molecules were docked to the corresponding protein binding pocket, and two representative examples (shown in blue and purple) are compared against the docked pose of the best-known active molecule.

For the pretraining part, we utilized 1.5 billion SAFE strings sampled from ZINC 22 by NovoMolGen study, which represents the largest pretraining dataset related to drug discovery for de novo molecule generation. A scaffold-based validation set and a random validation set were constructed to provide complementary perspectives on generalization.

### Overview of StructureSAFE

StructureSAFE is a structure-based molecular generative framework for both hit identification and lead optimization, which extends a pre-trained Llama-like autoregressive chemical language models with protein-ligand cross-attention adaptors (Figure 2). Given a protein pocket and an optional reference ligand, the model generates chemically valid molecules in SAFE string format. The system couples three components: (i) a dual-modality protein condition encoder, (ii) a deterministic ligand condition encoder, and (iii) a set of cross-attention adapters injected into a pretrained autoregressive backbone. Training combines a causal language modelling loss with a chemistry-aware Tanimoto correlation loss to finetune the ligand latent space, while classifier-free guidance (CFG) dropout enables conditional and unconditional generation from a single model.

### Base Chemical Language Model and Molecular Representation

During the training and inference, molecules are represented as SAFE strings, a line notation that encodes molecular graphs as ordered attachment-based fragment sequences. SAFE inherits SMILES grammar while imposing a BRICS fragmentation method that allows not only de novo hit generation but also multiple subtasks in lead optimization, including scaffold hopping, fragment elaboration and so on. A byte-pair encoding (BPE) tokenizer with a vocabulary of 500 tokens was trained on the SAFE-encoded training corpus took from NovoMolGen. Due to the limit of resources, stereochemistry information is taken away from SAFE strings and sequences are truncated or padded to a maximum length of 64 tokens (Figure 2B).

The backbone of our model is a Llama-architecture causal transformer with 300 million parameters: 32 transformer layers, hidden dimension d = 768, 12 attention heads, and feed-forward intermediate dimension 3,072. Flash Attention is used for all self-attention computations. The model is initialized from a pretrained checkpoint trained on a 1.5 billion unlabeled molecular corpus and subsequently fine-tuned with the conditioning framework described below.

### Dual-Modality Protein Condition Encoder

To give rich information about binding pocket and corresponding protein, protein structure and evolutionary context are encoded by combining two complementary representations: a Unimol pocket embedding ***x***_*pocket*_ ∈ ℝ^512^, capturing three-dimensional binding-site geometry, and an ESM-2 evolutionary embedding ***x***_*evo*_ ∈ ℝ^1280^, encoding sequence-derived conservation and co-evolutionary signals. For Unimol pocket embedding, only residues within 10 Å of reference ligand are considered, whereas the ESM-2 embedding is computed using the full protein sequence.

After generating the initial protein condition embeddings, both representations are projected into a shared latent space of dimension *C* = 512 via independent linear transformations, followed by layer normalization and dropout (p=0.1):

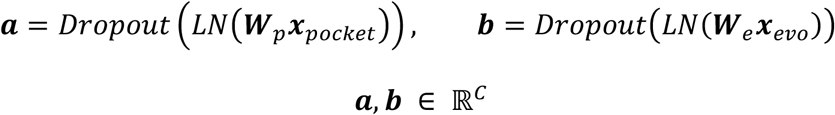

An early-fusion representation is constructed to integrate structural and evolutionary protein features by concatenating element-wise interactions and differences:

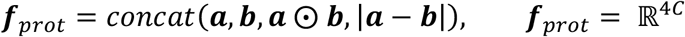

A two-layer multilayer perceptron (MLP) with GELU activations and hidden dimension max(512, 2*d*_*z*_) maps ***f***_*prot*_ to a protein condition embedding 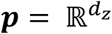, where *d*_*z*_ = 128:

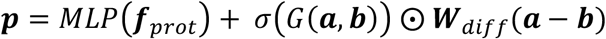

where *G*(·) denotes a two-layer gating network that produces a *d*_*z*_-dimensional modulation vector, *σ*(·)is the sigmoid function, and ***W***_*diff*_ is a learnable linear projection (Figure 2C). All linear layers are initialized using Xavier uniform initialization with zero biases.

### Ligand Condition Encoder

Ligand identity is encoded using two complementary representations: an interaction fingerprint (IFP) ***x***_*ifp*_ ∈ ℝ^16384^, capturing protein-ligand contact patterns at the binding site, and a Unimol molecular representation ***x***_*mol*_ ∈ ℝ^1536^, encoding both three-dimensional geometry and sequence-level information of the ligand. The ligand encoder is deterministic, produces a single condition vector ***µ*** in 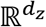 that summarizes the reference ligand condition for downstream conditioning.

The IFP tower compresses the high-dimensional fingerprint through three fully connected layers (16384 −> 512 −> 512 −> *d*_*embed*_, GELU activations, Dropout, and LayerNorm at each block) to final output ***a***. The molecular tower applies a two-layer MLP (1536 −> 512 −> *d*_*embed*_) to output ***b***. The resulting embeddings ***a*** and ***b*** are fused as following to produce the ligand molecular embedding ***µ***. Ligand molecular inputs are standardized using training-set statistics prior to encoding. We set *d*_*embed*_ = 256 in all experiments.

The two tower outputs **a** and **b** are fused using the same early-fusion strategy as for protein representations:

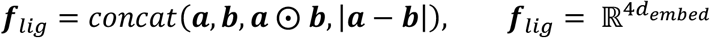

A gated residual fusion MLP maps ***f***_*lig*_ to a latent representation 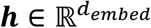, and a final linear readout head produces the ligand condition vector:

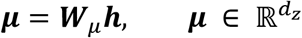

The vector mu is used directly as the ligand conditioning signal in the cross-attention layers (see Cross-Attention Conditioning) and as the input to the Tanimoto alignment loss (see Training objective, Figure 2C).

### Cross-Attention Conditioning

Protein and ligand condition signals are injected into the autoregressive backbone via lightweight cross-attention adapters inserted at all 32 transformer layers. Conditioning at every depth maximize the capacity of the cross-attention pathway to propagate the protein/ligand signal through the network.

The protein condition embedding p and ligand condition vector μ are independently expended into four condition tokens and projected into the transformer hidden space of dimension *d* = 768 via linear layers:

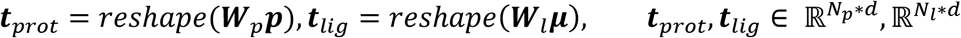

These are concatenated to form the conditioning matrix:

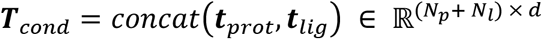

Decoupling the two modalities into separate tokens enables the cross-attention mechanism to independently regulate the contribution of protein structural information and ligand chemical identity at each layer. We use ***N***_*p*_ = ***N***_*l*_ = 4 in all experiments, giving a key-value memory of 8 tokens per sample. This richer K-V space allows cross-attention heads to selectively attend to different facets of the conditioning information (e.g. different sub-pocket regions or different ligand substructures) at each layer, while still being parameter-efficient compared with using the full encoded protein/ligand sequence.

At each conditioned layer *l*, the hidden states ***H***^(*l*)^ ∈ ℝ^*T* × *d*^ are updated via cross-attention. The layer-normalized hidden states serve as queries, while ***T***_*cond*_ provides keys and values:

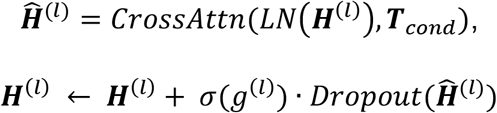

where *g*^(*l*)^ ∈ ℝ is a learnable scalar gate initialized to zero, such that *σ*(*g*^(*l*)^) = 0.5 at initialization. This design enables a smooth transition from the pretrained unconditional model, allowing the network to adaptively control the strength of conditioning. The cross-attention module uses the same number of attention heads (12) as the backbone self-attention (Figure 2B, 2C).

### Classifier-free guidance (CFG) dropout

To enable CFG at inference time, the ligand condition vector ***µ*** is randomlu replaced during training with a learnable null embedding 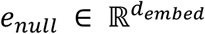 with probability ***p***_*drop*_ = 0.5. When dropped, ***t***_*lig*_ is computed from ***e***_*null*_ instead of ***µ***, while ***t***_*prot*_ is always retrained:

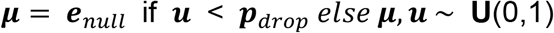

This single model therefore learns both the conditional distribution ***p***(***x***|*prot, lig*) and protein-only distribution ***p***(***x***|*prot*, ***e***_*null*_). At inference, the two predicted logits are combined as ***l***_*cfg*_ =***l***_*uncond*_ + *s* * (***l***_*cond*_ − ***l***_*uncond*_) with guidance scale *s* to amplify or decrease the ligand condition signal beyond what was directly trained.

### Training Objective

The total training loss consists of three components (Figure 2D):

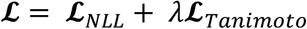

The negative log-likelihood term *L*_***N****LL*_ corresponds to the standard causal language modeling objective over the SAFE token sequence, with padding positions masked out. When CFG dropout is applied, the same NLL is computed with the null ligand embedding in place of ***µ***, so that the unconditional branch receives full language-modelling supervision.

To regularize the ligand latent space to be chemically meaningful, we introduce a Tanimoto correlation loss between pairwise distances in latent space and pairwise Tanimoto similarities of the corresponding molecules. For a training batch of B samples with ligand condition vectors ***µ***_*i*_ and pre-computed Morgan fingerprints ***fp***_*i*_ (radius 2, 2048 bits), define:

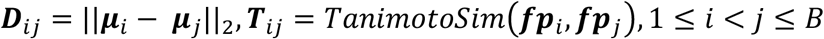

Let ***δ*** be the Pearson correlation coefficient between the upper-triangular entries of ***D*** and ***T***. The loss is defined as:

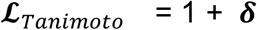

This loss reaches its minimum value of 0 when ***δ*** = −1, i.e. when Euclidean distances in the ligand latent space are perfectly anti-correlated with Tanimoto similarities (chemically similar ligands are mapped to nearby latent codes, and dissimilar ligands to distant ones). Batches with degenerate variance (***T***_*val*_ or ***D***_*val*_ below 1e-6) return zero loss to avoid numerical issues. *L*_*Tanimoto*_ operates on the deterministic ***µ*** and is unaffected by CFG dropout: every sample in a batch contributes its ***µ*** to the pairwise statistics, regardless of whether its conditioning was dropped on the NLL branch. The loss weight is *λ* = 0.1 in all reported experiments.

### Implementation Details

All experiments are conducted using the AdamW optimizer with *β*_1_ = 0.9, *β*_2_ = 0.95, weight decay of 0.01, and gradient clipping at a global norm of 5.0. The learning rate follows a cosine decay schedule with 10% linear warm-up, reaching a peak value of 5 × 10^−5^. Training is performed with a per-device batch size of 250 and 4 gradient accumulation steps (effective batch size of 1,000) for 10 epochs on the 5× augmented dataset. All computations use bfloat16 precision with TensorFloat-32 enabled for matrix multiplications. The model is implemented in PyTorch using the Hugging Face Transformers framework with custom data modules. Training is monitored using Weights & Biases. The random seed is fixed to 2025 for reproducibility. We perform full end-to-end fine-tuning of all model parameters, resulting in approximately 330 million trainable parameters.

### Inference Details

Depending on the availability of ligand information, inference is performed under two settings: pocket-conditioned generation and ligand-aware and pocket-conditioned generation. In the pocket-conditioned setting, the guidance scale *s* is set to 1 with the null ligand embedding ***e***_*null*_ are feed to the network. In the ligand-aware setting, the normal calculated ***µ*** are feed to the network with *s* set to 1.5 to amplify the ligand condition. The generated SAFE strings are converted to SMILES strings as the final molecular outputs. For lead optimization tasks, the regions of a molecule to be modified are specified using dummy atoms in the SMILES representation. The modified SMILES is converted to SAFE format and tokenized as model input. Further details of the SAFE representation are provided in the original work.

### Baselines

Since bdnv2 is a relatively new dataset and we did the data splitting by ourselves, it is relatively hard to directly compare the performance between our model and other people’s model. Thus, to allow better comparison, we utilized CrossDocked2020 as training set to finetune our model (refer to as StructureSAFE-c), which allows direct comparison with all other models trained using same dataset. We utilized MolGenBench benchmarking dataset,^21^ generating 1,000 molecules for each of 120 target proteins, resulting in a total of 120,000 molecules for metrics calculation. After calculating metrics using MolGenBench, the result is mainly compared with other models evaluated in MolGenBench study without running the generation using those models by ourselves.

### Evaluation metrics

We evaluated *de novo* generation and lead optimization performance of our model on both test set and MolGenBench dataset using following metrics:

- **Vina score**. The Vina score estimates the binding affinity between ligands and the receptor. We applied the python version of AutoDock Vina v1.2.7 for calculating all Vina scores and prepared protein and ligands using ‘scrub.py’, ‘mk_prepare_ligand.py’ and ‘mk_prepare_recepor.py’. After docking, the best vina score for each ligand are recorded for calculating statistics, with more negative values indicate better predicted binding affinity.
- **QED**. The quantitative estimate of drug-likeness (QED) estimates the potential of a small molecules to become an oral drug based on the property distributions of approved drug molecules. The score ranges from 0 to 1, with higher values indicating better druglikeness.
- **SA**. Synthetic accessibility (SA) score measures the ease of synthesizing a molecule, based on fragment frequency distributions from known compounds and structural complexity penalties. The original score ranges from 1 to 10, and we normalized it to 0 to 1, with higher values indicating easier synthesis.
- **Ring_freq > 100**.^12, 22^ This metric measures the percentage of ring systems in the generated molecules that occur more than 100 times in the ChEMBL ring system dataset. The value ranges from 0% to 100%, with higher values indicating better chemical plausibility and drug-likeness.

## Result

### StructureSAFE Model Development

StructureSAFE is a Llama-based chemical language model conditioned on protein pocket information and optional ligand information, designed for structure-based hit generation and lead optimization. The model architecture and sampling process are illustrated in Figure 2. The backbone of StructureSAFE is a decoder-only Transformer based on Llama architecture adopted from NovoMolGen model family with 300M parameters (Method).^13^ To learn the grammar of SAFE strings and enhance the chemical plausibility of generated molecules, the model was pretrained on 1.5 billion molecules from the NovoMolGen dataset, enabling it to internalize rich chemical prior information. This pretraining corpus represents the largest dataset employed for *de novo* generation pretraining using the SAFE string representation. Following pretraining, we generated 10,000 molecules and evaluated a set of relative performance metrics to assess the generative capability of the backbone model. As shown in Table S1, nearly all generated molecules are valid (validity = 0.999) with no duplicates, demonstrating that the backbone successfully learned chemical prior information from the training data and can reliably produce valid and unique molecules. Furthermore, the internal diversity score (0.848) and medicinal chemistry filter pass rate (0.854) are both satisfactory, indicating that the backbone captures the underlying training data distribution and is capable of generating structurally diverse, drug-like molecules (Methods).

With confidence established in our backbone model, we proceeded to the fine-tuning stage. To enable structure-based molecular generation and optimization, the model requires fine-grained structural information of the target binding pocket. To this end, we employed the pretrained UniMol model^23^ as our pocket encoder to capture detailed three-dimensional pocket structural information. Additionally, given that evolutionary information has been shown to facilitate structure prediction and protein–ligand interaction modeling in AlphaFold^4^ and the Chai^24, 25^ model series, we incorporated ESM-2^26^ to encode sequence-level and evolutionary information. The structural and evolutionary representations are subsequently concatenated and passed to the model as a unified protein context. In practical hit identification and lead optimization scenarios, researchers may already have access to known ligands that bind to the target of interest. In such cases, it is desirable to leverage these known ligands as templates to guide de novo generation or optimization toward higher-affinity candidates. To accommodate this, we introduced a ligand module that encodes ligand information using UniMol embeddings and captures protein–ligand interaction features via PLEC interaction fingerprints.^27^ Classifier-Free Guidance (CFG) is incorporated to allow ligand information to serve as an optional conditioning input. The final representations from the ligand module and protein module are combined and used to condition the generation process via cross-attention (Figure 2).

To prevent data leakage and evaluate the generalizability of the fine-tuned model, we partitioned the BindingNet v2 dataset (comprising 689,796 protein–ligand complexes in total) based on both protein sequence similarity and scaffold overlap between the training and test sets (Methods). Following this stratified splitting procedure, 228 proteins and their corresponding ligands were selected as the test set. The remaining 688,102 protein–ligand complexes were retained as the training set, which was further augmented by randomizing SAFE strings five times per sample, yielding a total of 3,440,510 training pairs (Figure 2A).

### Model Performance on *De Novo* Hit Identification

Following fine-tuning, the performance of StructureSAFE on hit identification was comprehensively evaluated using both the test set and the MolGenBench dataset. For the test set evaluation, StructureSAFE was applied to generate 50 ligands per protein, yielding a total of 11,400 molecules. For protein-only conditioned generation, as shown in Table 1, the generated molecules exhibit an average synthetic accessibility (SA) score of 2.92 and an average quantitative estimate of drug-likeness (QED) of 0.54, reflecting favorable drug-like properties across the generated set. For reference, the true ligands in the test set have average SA and QED values of 3.22 and 0.49, respectively, while ligands in the training set have corresponding values of 3.13 and 0.51. The generated molecules also demonstrate high uniqueness, with 11,312 out of 11,400 molecules (99.2%) being non-redundant, underscoring StructureSAFE’s strong capacity for generating structurally diverse compounds.

**Table 1.** Performance of StructureSAFE under two conditioning modes on the test set. Detailed definitions and explanations of each evaluation metric are provided in the Methods section. “Average best-affinity score” denotes the mean Vina docking score of the top-ranked generated molecule per target, averaged across all targets.

|  | Test set | Generated molecules<br>(Protein-only condition) | Generated molecules<br>(Protein & ligand<br>condition) |
| --- | --- | --- | --- |
| QED (↑) | 0.49 | 0.54 | 0.47 |
| SA (↓) | 3.22 | 2.92 | 4.37 |
| Vina docking score (Average)<br>(↓) | -8.45 | -7.82 | -6.85 |
| Vina docking score (Average<br>best_affinity) (↓) | N/A | -10.00 | -9.02 |
| Ring_freq > 100 (%) (↑) | 76.75% | 76.83% | 44.67% |

To further assess binding affinity, we docked the generated molecules to their corresponding protein pockets and compared the resulting docking scores against those of the best-known binders for each target. As shown in Table 1, while the average docking score of generated molecules across all targets is lower than that of the best-known binders (−7.82 vs. −8.45), a substantial proportion of generated molecules are predicted to exhibit superior binding affinities, with an average best-affinity score of −10.00. This finding highlights the potential of StructureSAFE to generate high-affinity binders for previously unseen targets. From a medicinal chemistry perspective, 76.83% of generated molecules contain ring systems with a frequency greater than 100 (Ring_freq > 100), a rate comparable to that observed in the test set ligands, indicating that the model has successfully internalized chemical prior information from the training data and reliably generates chemically plausible structures.

Beyond protein-only conditioned generation, ligand information can additionally be incorporated to condition StructureSAFE via classifier-free guidance (CFG). We therefore evaluated generation performance under dual conditioning, using both protein information and ligand information from the test set at a CFG scale of 1.5, which modestly amplifies the contribution of ligand information to the generation process. As shown in Table 1, molecules generated under dual conditioning exhibit an average QED of 0.47 and an average SA score of 4.37, with a mean docking score of −6.85 and an average best-affinity score of −9.02 per target. The substantial shift in QED and SA statistics relative to protein-only generation suggests that the two conditioning modes explore distinct regions of chemical space. To investigate this further, we computed the nearest-neighbor Tanimoto similarity (NNTS) between molecules generated under protein-only conditioning and those generated under dual conditioning. As shown in Table S1, the mean NNTS between the two groups is 0.28, with only 24.2% of molecules exceeding an NNTS of 0.3 (empirically established threshold for structural similarity). Taking together, these results, in conjunction with the divergent QED and SA profiles, confirm a substantial shift in the chemical space explored by the two generation modes.

Given that BindingNet v2 is a relatively recent dataset that has seen limited adoption for model training, we conducted a fair comparison with existing models using the MolGenBench benchmark, with CrossDocked2020 as the training set.^28^ We retained the five-fold data augmentation strategy and all hyperparameters, with the exception of the number of training epochs and learning rate, which were adjusted to account for the approximately seven-fold reduction in training data size. The model trained on the CrossDocked2020 dataset is hereafter referred to as StructureSAFE-c. As shown in Table 2, StructureSAFE-c achieves strong performance across all evaluated metrics relative to state-of-the-art models on the MolGenBench dataset, ranking in the top 3 for the majority of metrics with the exception of Diversity. Notably, StructureSAFE-c outperforms most competing models by a substantial margin on the Ring_freq > 100 metric, only chemical language model TamGen and the flow-matching model PocketFlow showed close performance. This result highlights the effectiveness of our pretraining and fine-tuning strategies in capturing chemical prior information and generating chemically plausible molecules. With respect to the re-discovery of known active molecules and scaffolds (Figures 3A and 3B), StructureSAFE-c demonstrates competitive performance among top-ranked models, achieving second-place performance in regenerating known SMILES for proteins within CrossDocked2020, and first-place performance in regenerating known scaffolds for proteins outside of CrossDocked2020.

**Table 2.** Benchmarking of StructureSAFE-c against SOTA structure-based generative models on the CrossDocked2020 test set. We applied the same metrics as MolGenBench study,^21^ and all performance values for models other than StructureSAFE-c are sourced directly from the MolGenBench benchmark report.

|  | PocketFlow | TamGen | MolCraft | Pocket2Mol | DiffSB DD-C | DiffSB DD-M | TargetDiff | DecompDiff | SurfGen | FLAG | StructureSAFE-c |
| --- | --- | --- | --- | --- | --- | --- | --- | --- | --- | --- | --- |
| Validity (↑) | 1.00 | 1.00 | 0.96 | 0.88 | 0.78 | 0.72 | 0.68 | 0.59 | 0.37 | 0.14 | 1.00 |
| QED (↑) | 0.48 | 0.57 | 0.50 | 0.44 | 0.50 | 0.46 | 0.32 | 0.52 | 0.38 | 0.59 | 0.54 |
| SA (↑) | 0.78 | 0.76 | 0.65 | 0.66 | 0.60 | 0.61 | 0.54 | 0.58 | 0.61 | 0.64 | 0.74 |
| Uniqueness (↑) | 0.92 | 0.27 | 1.00 | 1.00 | 1.00 | 1.00 | 1.00 | 0.93 | 1.00 | 1.00 | 1.00 |
| Diversity (↑) | 0.88 | 0.84 | 0.88 | 0.83 | 0.90 | 0.90 | 0.89 | 0.84 | 0.81 | 0.90 | 0.87 |
| Ring_freq > 100 (↑) | 0.74 | 0.75 | 0.26 | 0.27 | 0.31 | 0.31 | 0.24 | 0.42 | 0.06 | 0.56 | 0.79 |

**Figure 3.**
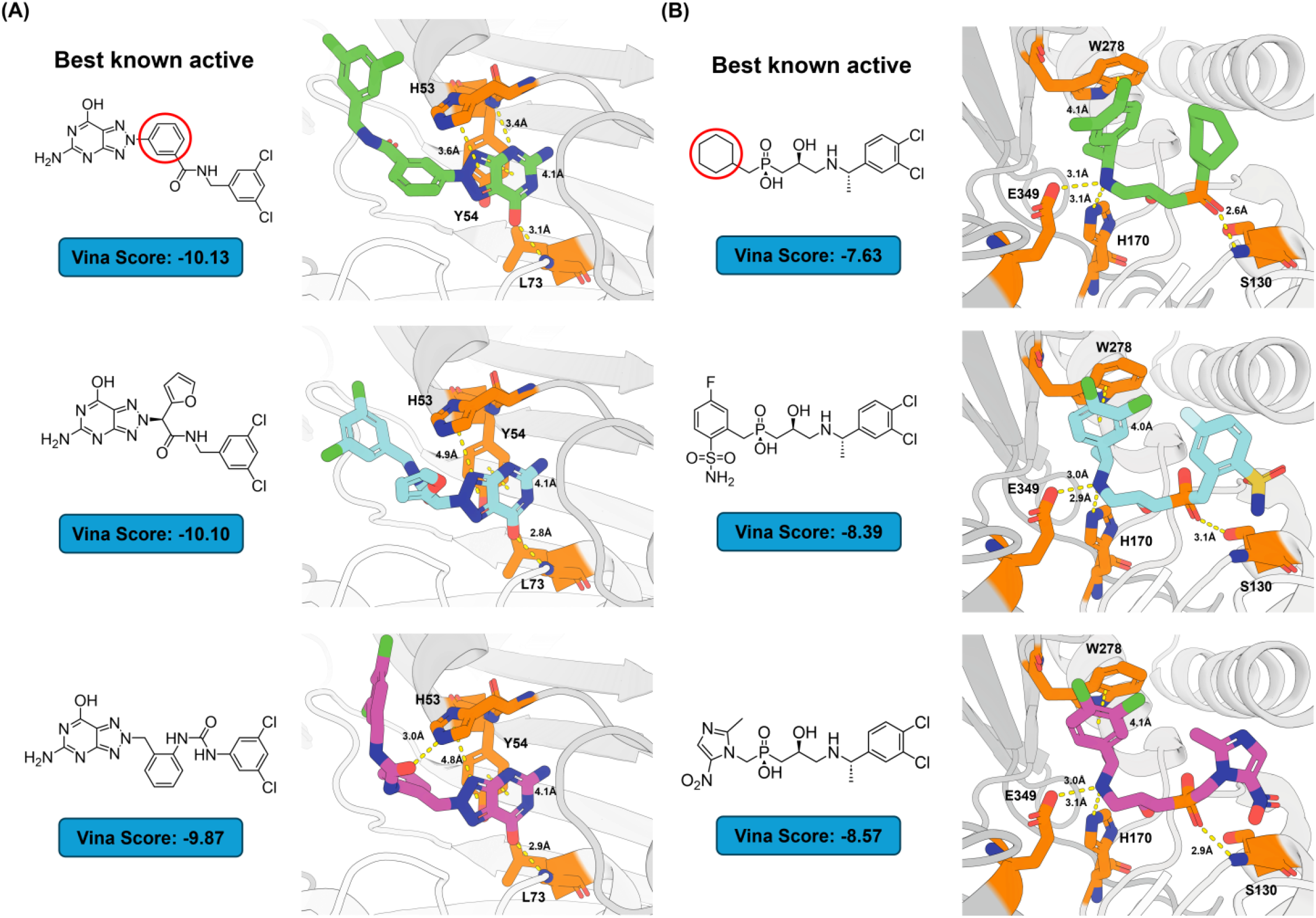
In silico lead optimization case studies of StructureSAFE on DHNA and the GABA-B receptor. Protein–ligand interactions are indicated by yellow dashed lines, and fragments circled in red denote the substitution sites targeted for replacement. (A) Application of StructureSAFE to the antimicrobial target DHNA for a scaffold hopping subtask. Generated molecules are compared against the best-known active ligand (ChEMBL3217378, IC_50_ = 68 nM) with respect to predicted binding interactions and Vina docking scores. (B) Application of StructureSAFE to the GABA-B receptor for a fragment elaboration subtask. Generated molecules are compared against the best-known active ligand (ChEMBL2391908, Ki = 2.9 nM) with respect to predicted binding interactions and Vina docking scores.

To further demonstrate the *de novo* hit identification capability of StructureSAFE, we selected two therapeutically relevant targets that are not in our training set for case study analysis: Relaxin-3 receptor 2 (RXFP4, a promising GPCR target implicated in immuneandmetabolicdiseases)^29^andUDP-N-acetylmuramoylalanine-D-glutamyl-lysine-D-alanyl-D-alanine ligase (MurF, a potential antibacterial enzymatic target)^30^. Following our standard evaluation protocol, we generated 50 molecules per target under each conditioning mode (protein-only and protein–ligand), yielding 100 candidate molecules per target in total. As shown in Figure 3C, the best-known ligand for MurF (ChEMBL272818, IC_50_ = 22.0 nM) achieves a docking score of −10.96. Analysis of its binding interactions reveals that the sulfonamide group forms a hydrogen bond with Asn328, the nitrile group forms two hydrogen bonds with Ala48 and Arg49, and the phenol group forms a hydrogen bond with the backbone oxygen of Leu360. These interactions are collectively expected to contribute to the high binding affinity of this known ligand.

Among the generated molecules, the first candidate (ligand in blue, docking score = −11.09) retains the sulfonamide group present in the best-known ligand and similarly forms a hydrogen bond with Asn328, recapitulating one of the key interactions of the reference compound. In addition, it establishes a novel hydrogen bond with Phe31, likely contributing to its favorable predicted docking score. The second molecule (ligand in purple, docking score = −12.56) adopts a more lipophilic binding mode, with hydrophobic interactions serving as the primary driver of its predicted affinity. While the lipophilic character of this molecule may introduce risks of off-target effects and metabolic liabilities, these properties could be systematically addressed through subsequent lead optimization once bindingaffinityisexperimentallyconfirmed.

For RXFP4, as shown in Figure 3D, the best-known ligand (ChEMBL5094669, IC_50_= 0.9 nM, docking score = −8.57) forms three hydrogen bonds with residues Tyr204, Arg208, and His263. In addition, the positively charged Arg208 engages in a cation–π interaction with the pyridine ring of the ligand, further enhancing its predicted binding affinity. Among the generated molecules, the first candidate (docking score = −10.8) similarly forms a cation–π interaction with Arg208, recapitulating this key interaction observed in the best-known ligand. It additionally engages in two π–π stacking interactions with Trp97 and His299, respectively, as well as a hydrogen bond with Asn262. The second generated molecule (docking score = −10.09) forms three hydrogen bonds with residues Arg194, Tyr204, and Arg208, along with a π–π stacking interaction with Trp97. Taken together, both generated molecules engage Arg208, which is a key residue also targeted by the best-known ligand, and exhibit relatively balanced lipophilicity profiles, rendering them suitable candidates for further optimization and development.

Overall, StructureSAFE’s performance on *de novo* generation demonstrates its capacity to produce drug-like, potentially high-affinity hit compounds given protein structural information alone, while exploring distinct regions of chemical space when ligand information is additionally provided as conditioning input.

### Model Performance on Lead Optimization

To evaluate StructureSAFE’s performance on lead optimization, we randomly removed one fragment from the SAFE string of each ground-truth ligand in the test set and prompted the model to generate 50 novel fragment replacements per target–molecule pair under both conditioning modes. This protocol collectively covers a broad range of lead optimization subtasks, including but not limited to linker design, fragment elaboration, R-group enumeration, and scaffold hopping. For protein-only conditioned generation, the generated molecules exhibit average SA and QED values of 3.61 and 0.42, respectively. Relative to *de novo* generation, the average docking score improves modestly, likely because the majority of the parent scaffold is retained during fragment replacement (Table 3). Nevertheless, the average docking score of generated molecules does not surpass that of the best-known binders (−8.08 vs. −8.45). Despite this, a considerable proportion of generated molecules are predicted to exhibit superior binding affinities, with an average best-affinity score of −9.43. These higher-affinity fragment replacements could serve as a prioritized candidate pool for downstream high-precision calculations such as free energy perturbation (FEP) or experimental synthesis following inspection and filtering by medicinal chemists.

**Table 3.** Performance of StructureSAFE under two conditioning modes on the test set for lead optimization tasks. Detailed definitions and explanations of each evaluation metric are provided in the Methods section. “Average best-affinity score” denotes the mean Vina docking score of the top-ranked generated molecule per target, averaged across all targets.

|  | Test set | Generated molecules<br>(Protein-only condition) | Generated molecules<br>(Protein & ligand<br>condition) |
| --- | --- | --- | --- |
| QED (↑) | 0.49 | 0.42 | 0.41 |
| SA (↓) | 3.22 | 3.61 | 3.88 |
| Vina docking score (Average)<br>(↓) | -8.45 | -8.08 | -7.84 |
| Vina docking score (Average<br>best_affinity) (↓) | N/A | -9.43 | -9.23 |
| Ring_freq > 100 (%) (↑) | 76.75% | 62.52% | 59.71% |

When ligand information from the best-known binders is additionally incorporated as a conditioning input (CFG scale = 1.5), the generated molecules exhibit average SA and QED values of 3.88 and 0.41, respectively, with a mean docking score of −7.84 and an average best-affinity score of −9.23, further demonstrating the potential of StructureSAFE to propose high-affinity lead optimization vectors. However, examination of the Ring_freq > 100 metric reveals a more nuanced picture: while dual conditioning yields an approximately 15% improvement over de novo generation, protein-only conditioned lead optimization shows a substantial performance decrease on this metric. Compared with the corresponding values for the test set ligands, this observation suggests that capturing chemical prior information is more challenging in the context of lead optimization, likely owing to the inherently greater complexity lead optimization subtasks relative to *de novo* generation.

To evaluate StructureSAFE’s performance on target-specific lead optimization in silico, we selected two additional receptors that are not in training set as case studies: Dihydroneopterin aldolase (DHNA, an antimicrobial enzymatic target)^31^ and the gamma-aminobutyric acid type B (GABA-B) receptor (a GPCR target implicated in neurological and metabolic diseases).^32^ For DHNA, as shown in Figure 4A, the best-known ligand (ChEMBL3217378, IC_50_ = 68 nM, docking score =−10.13) establishes two hydrogen bonds with His53 and Leu73, and additionally engages in two π– π stacking interactions with His53 and Tyr54. These interactions collectively account for the high predicted docking score of this ligand. We applied StructureSAFE to perform a scaffold-hopping task on this ligand, replacing the central benzene ring while retaining the remainder of the scaffold, conditioned solely on protein structural information, with the aim of improving binding affinity. The first generated molecule also forms π–π stackings with His53 and Tyr54, with similar hydrogen bond on Leu73 while positioning a furan ring in the solvent exposed region. The second generated molecule similarly forms π–π stacking interaction and also a novel hydrogen bond with His53, meanwhile keeping other interactions with Leu73 and Tyr54. However, both generated molecules are predicted to disrupt the hydrogen bond observed in the best-known ligand with backbone nitrogen of Tyr54, which may account for their suboptimal docking scores. It is worth noting that the docking protocol employs a rigid receptor approximation; in reality, the receptor may undergo subtle conformational adjustments to accommodate and preserve favorable hydrogen bonding interactions. We therefore consider these designs to remain worthy of experimental investigation. Furthermore, both generated molecules exhibit an increased fraction of sp^3^ carbon atoms, conferring greater three-dimensional character and conformational flexibility, which is generally favored in medicinal chemistry for potential better affinity and metabolic profiles.

For the GABA-B receptor, the best-known ligand (ChEMBL2391908, Ki = 2.9 nM, docking score = −7.63) forms two hydrogen bonds with Ser130 and His170, a π –π stacking interaction with Trp278, and a salt bridge with Glu349 (Figure 4B). To further improve predicted binding affinity, we applied StructureSAFE to replace the solvent-exposed cyclohexane ring of this ligand with alternative fragments (fragment elaboration task). The first generated molecule (docking score = −8.39) introduces a novel hydrogen bond with Ser130 while preserving the majority of interactions observed in the best-known ligand with the remaining binding site residues. The second generated molecule (docking score = −8.57) almost copied the interactions of reference ligand with slight change of binding conformation. The overall improvement in predicted docking scores for both candidates likely arises from the introduction of additional polar groups into the solvent-exposed region, which may contribute favorable desolvation energy.

Taken together, these two case studies demonstrate that StructureSAFE can be flexibly applied across distinct lead optimization subtasks and is capable of generating structurally diverse, chemically meaningful candidates for consideration by medicinal chemists in subsequent synthesis and experimental validation.

## Discussion

In our previous work, we demonstrated that chemical plausibility represents a critical liability for structure-based generative models, and hypothesized that pretraining and data augmentation strategies could substantially improve performance in this regard.^12^ Chemical language models are particularly well-positioned to benefit from such strategies, given the relative ease with which large-scale pretraining and task-specific fine-tuning can be incorporated into their training pipelines. In the present study, we observe that two chemical language models, StructureSAFE and TamGen^16^, consistently outperform the majority of graph-based models lacking pretraining in generating chemically plausible molecules. A notable exception among graph-based models is PocketFlow^33^, whose comparatively strong chemical plausibility performance likely stems from its pretraining on molecules from ZINC prior to fine-tuning on the CrossDocked2020 dataset. Collectively, these results underscore the importance of capturing chemical prior information during model development, whether through large-scale pretraining, data augmentation, or fragment-based generation strategies.^12^

During training, we observed that some trained models in an overfitted regime (i.e. almost ignore protein/ligand conditioning, simply generating ligands randomly) paradoxically achieved equal or superior performance on several metrics, including QED, SA, validity, and diversity (Table S3). The same phenomenon is also observed for TamGen.^21^ Even on protein-relevant metrics such as molecular docking scores, performance remained comparable, consistent with observations reported elsewhere that structure-based generative models can hardly produce molecules that match or outperform randomly sampled ChEMBL compounds on docking-based evaluation metrics.^34^ While an unconditional, high-quality molecular generator may still hold practical utility for large-scale virtual screening, this observation highlights a fundamental limitation of current evaluation frameworks and underscores the pressing need for more discriminative and practically meaningful metrics for assessing the true performance of structure-based generative models.

Beyond evaluation metrics, the choice of fragmentation method also exerts a substantial influence on the applicable domains of fragment-based generative models. The BRICS fragmentation scheme employed in SAFE string construction tends to produce relatively large fragments that are derived from a limited set of retrosynthetic rules and bond-breaking patterns. This restricted coverage of fragmentation methods may in turn constrain the structural diversity of generated molecules and introduce systematic biases toward specific reaction types. More sophisticated fragmentation strategies like learned fragmentation approaches in which a model is trained to identify chemically meaningful breaking points hold promise for broadening the scope and improving the quality of fragment-based molecular generation.^35^

Looking ahead, the generative performance of StructureSAFE could be further enhanced by incorporating reinforcement learning to guide molecule generation toward compounds satisfying specific multi-parameter optimization criteria. Additionally, extending the model to support direct 3D molecular generation would enable more explicit utilization of the rich three-dimensional structural information encoded in protein binding pockets, albeit at the cost of reduced pretraining data availability. In summary, StructureSAFE is a ready-to-use, versatile platform that can provide high-quality candidate molecules to support medicinal chemists and complement rigorous physics-based methods in the selection of final candidates for synthesis and biological validation.

## Summary

In this study, we developed StructureSAFE, a structure-aware chemical language model capable of generating high-quality candidate molecules conditioned on protein structural information and, optionally, known ligand information, for both de novo hit identification and lead optimization tasks. Comprehensive evaluation on the test set and benchmarking against the MolGenBench dataset demonstrates that StructureSAFE achieves state-of-the-art performance in generating diverse, drug-like, and synthetically accessible molecules, while maintaining target relevance as evidenced by competitive docking scores and successful regeneration of known ligands. Notably, compared to the majority of graph-based generative models, StructureSAFE achieves a substantial improvement in the chemical plausibility of generated molecules, reflecting its effective internalization of chemical prior information during pretraining. In silico case studies across four therapeutically relevant targets further validate StructureSAFE’s capacity to generalize across both hit identification and lead optimization tasks, positioning it as one of the few structure-based generative models capable of unifying these two complementary objectives within a single framework. The generated molecules not only recapitulate key interactions formed by high-affinity reference ligands but also introduce novel interactions with the potential to further enhance binding affinity. In summary, StructureSAFE represents a versatile and practical tool that can generate candidates for both computational scientists further filtering and medicinal chemists’ selection and synthesis in real-world hit identification and lead optimization campaigns.

